# Long-read metagenomics of soil communities reveals phylum-specific secondary metabolite dynamics

**DOI:** 10.1101/2021.01.23.426502

**Authors:** Marc W. Van Goethem, Andrew R. Osborn, Benjamin P. Bowen, Peter F. Andeer, Tami L. Swenson, Alicia Clum, Robert Riley, Guifen He, Maxim Koriabine, Laura Sandor, Mi Yan, Chris G. Daum, Yuko Yoshinaga, Thulani P. Makhalanyane, Ferran Garcia-Pichel, Axel Visel, Len A. Pennacchio, Ronan C. O’Malley, Trent R. Northen

**Affiliations:** Molecular EcoSystems Biology Division, Lawrence Berkeley National Laboratory, 1 Cyclotron Rd, Berkeley, CA, 94720, USA; DOE Joint Genome Institute, Lawrence Berkeley National Laboratory, 1 Cyclotron Rd, Berkeley, CA, 94720, USA; Centre for Microbial Ecology and Genomics, Department of Biochemistry, Genomics and Microbiology, University of Pretoria, Lynnwood Rd, Hatfield. Pretoria, 0028, South Africa; Center for Fundamental and Applied Microbiomics, Biodesign Institute, Arizona State University, Tempe, Arizona, USA; School of Life Sciences, Arizona State University, Tempe, Arizona, USA

**Keywords:** Long-read metagenomics, secondary metabolism, metatranscriptomics, biological soil crust, soil microbiome

## Abstract

Microbial biosynthetic gene clusters (BGCs) encoding secondary metabolites are thought to impact a plethora of biologically mediated environmental processes, yet their discovery and functional characterization in natural microbiomes remains challenging. Here we describe deep long-read sequencing and assembly of metagenomes from biological soil crusts, a group of soil communities that are rich in BGCs. Taking advantage of the unusually long assemblies produced by this approach, we recovered nearly 3,000 BGCs for analysis, including 695 novel, full-length BGCs. Functional exploration through metatranscriptome analysis of a 3-day wetting experiment uncovered phylum-specific BGC expression upon activation from dormancy, elucidating distinct roles and complex phylogenetic and temporal dynamics in wetting processes. For example, a pronounced increase in BGC transcription occurs at night in cyanobacteria but not in other phyla, implicating BGCs in nutrient scavenging roles and niche competition. Taken together, our results demonstrate that long-read metagenomic sequencing combined with metatranscriptomic analysis provides a direct view into the functional dynamics of BGCs in environmental processes and suggests a central role of secondary metabolites in maintaining phylogenetically conserved niches within biocrusts.

---

A fundamental challenge in understanding the ecological functions of secondary metabolites (also known as specialized metabolites or natural products) is that most biosynthetic gene clusters (BGCs) are harbored by uncultivated microbes and require specific native contexts for activation^1^. The majority of BGCs encoding secondary metabolites are not usually expressed under standard cultivation conditions in the laboratory^2^ and their products have therefore been termed ‘secondary’ metabolites. A universal feature of BGCs is their modular, co-localized gene architecture^3^ and large size, frequently spanning tens of thousands of base pairs. Bacterial secondary metabolites play critical ecological roles in mediating communication, antagonistic interactions, nutrient scavenging, and have historically been a primary source for antibiotic drug development^4^; in fact more than half of registered drugs are based on natural secondary metabolites^5^. Additionally, secondary metabolites have applications in agriculture^6^, biomaterials^7^, biofuels^8^, and cosmetics^9^.

Previous work has demonstrated the potential for deep “shotgun” metagenomic sequencing to directly characterize BGCs from environmental samples^10 11^, but the assembly of full-length BGCs from short reads is associated with significant limitations^12^. Alternative techniques include the use of clone libraries^1^ or innovative sequence-based analyses^13, 14^ including the reconstruction of uncultivated microbes as metagenome-assembled genomes (MAGs; reviewed in^15^). However these approaches typically only give access to dominant members of the community, while often omitting members of the ‘rare biosphere’^16^.

We also know remarkably little about the transcription of BGCs in nature or how the environment regulates their production^17^ especially in soils. This information is critical in understanding how often secondary metabolites are produced in natural communities. Biological soil crusts (biocrusts) are the world’s most extensive biofilms and together cover up to 12% of total soil surface area^18^. Initial studies have suggested that they are rich in secondary metabolites^19^. *Cyanobacteria* dominate biocrust communities, specifically *Microcoleus* spp. that drive biocrust establishment by stabilizing the soil surface, both preventing erosion and improving soil fertility through the release of photosynthate^20, 21^. In contrast to many other types of soil environments, biocrusts are easily transferable to the laboratory, which allows for controlled interrogation of relevant environmental processes such as wetting dynamics. In native environments, rain events suspend microbial dormancy in biocrust and cause dramatic shifts to community structure and both primary and secondary metabolite release^22^. The secondary metabolites produced by microbes upon wetting are known to include antimicrobial compounds thought to provide a selective advantage^23^, yet the majority of secondary metabolites encoded in the genomes of biocrust community members remain unidentified^24^. *Cyanobacteria* are known secondary metabolite producers^3, 25^ but most studies have focused on aquatic cyanobacteria, leaving the secondary metabolites of terrestrial cyanobacteria largely underexplored^26, 27^.

We combined long- and short-read metagenomic sequencing to produce ultra-large assemblies that enabled BGC discovery. We then mapped time-series metatranscriptomes to gain insight into the environmental cues governing BGC expression in biocrusts. Our results showed that thousands of gene clusters could be extracted from assembled long-read metagenomes which gave insight into the secondary metabolism of both rare and dominant microbial taxa. Coupling these results to metatranscriptomics indicated that most BGCs were transcribed after a simulated rain event, and that cyanobacteria dominated secondary metabolism.

## Long-read sequencing permits access to ultra-long gene clusters

Biocrust samples were collected from Moab, UT, USA (Fig. 1a), and transported to the JGI in petri dishes that maintain the physical structure of the crust. We then extracted and prepared high-molecular-weight DNA was extracted and prepared from intact biocrust samples for both long- and short-read metagenomic sequencing (Fig. S1). In total, we sequenced eight SMRT cells from three libraries yielding 156.3 Gb from 36.7 million reads, where half of all sequenced bases were contained in reads of 5 kb or longer, while the longest read was 167 kb. The average read length was 3,084 bp while the mean *N50* value was 4,070 bp. Both statistics were augmented by the Sequel II library which comprised 108 Gb of sequence in just 19.1 million reads.

**Fig 1.**
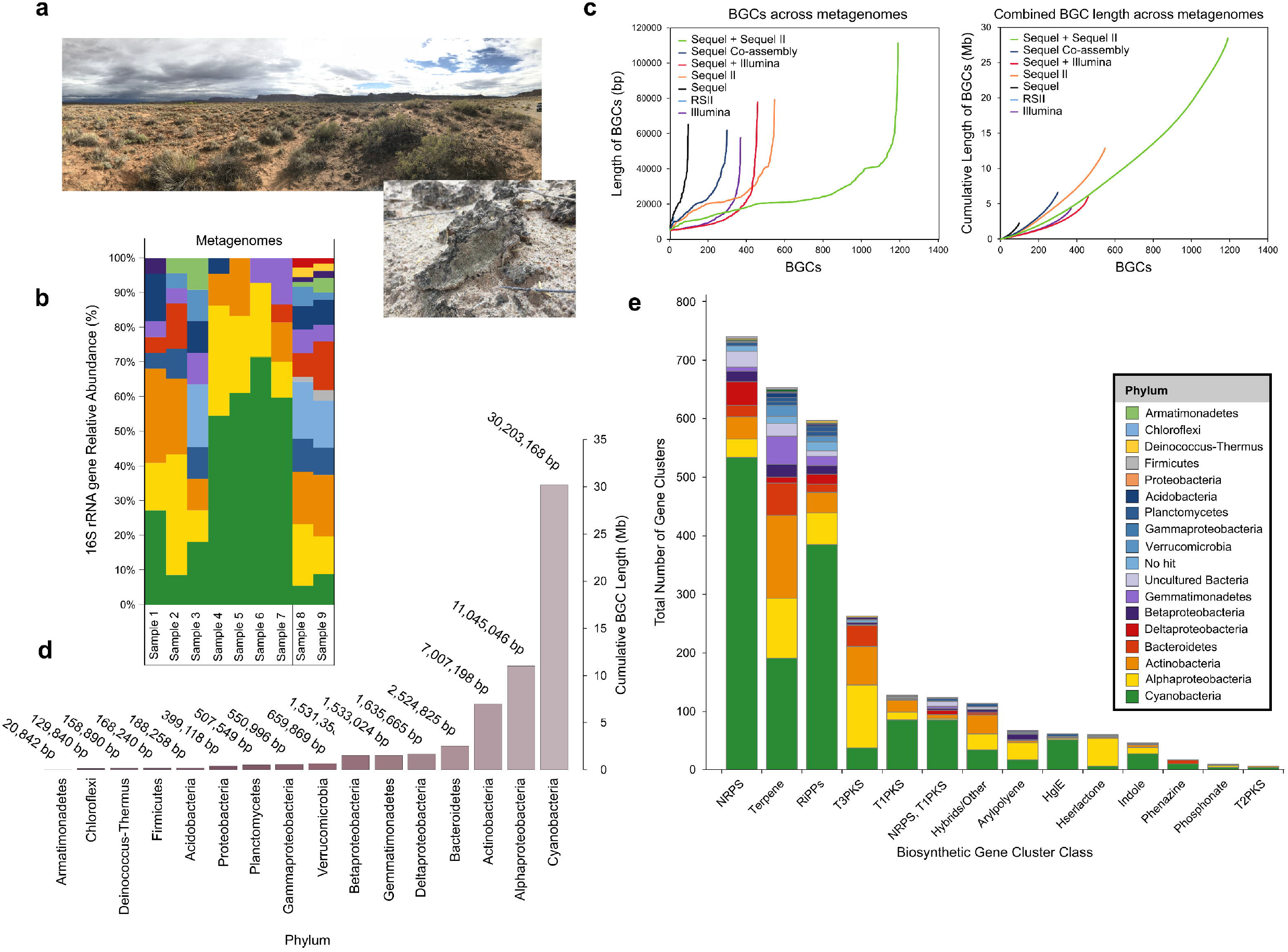
Secondary metabolism of biocrust. **a**, Field sampling location in the Green Butte Site near Canyonlands National Park (Moab, UT) with the biological soil crust inlay showing the characteristic green coloration. **b**, Taxonomic composition of the metagenomes based on exact sequence variants (ESVs) of 16S rRNA genes across sequencing platforms. Relative abundances were calculated after assigning taxonomy against the SILVA reference database. **c**, Left panel shows the number of Biosynthetic Gene Clusters (BGCs) recovered from each assembly, arranged from shortest to longest. Right panel shows the cumulative length of BGCs recovered from each metagenome in Megabases (Mb). **d**, Taxonomic distribution of BGCs in megabase pairs (Mb) at the phylum or class level. **e**, BGCs longer than 5 kb from each major class of secondary metabolism, colored by putative phylum-level assignments.

The 2 short-read Illumina libraries provided an additional 20 Gb of sequence (Table S1). To obtain an initial phylogenetic profile of the communities under investigation we performed full-length 16S rRNA gene analysis using exact sequence variants (ESVs) which showed that *Cyanobacteria*, and particularly *Microcoleus vaginatus*, were dominant biocrust community members, with major representations of *Actinobacteria* and *Alphaproteobacteria* (Fig. 1b) which is generally consistent with the known community composition of these biocrusts^28^. Overall, the biocrust are less complex than other desert soil communities^29^ yet are notably richer in cyanobacteria.

To access biosynthetic gene clusters, we individually assembled the biocrust metagenomes into contiguous sequences (contigs). Using both Canu^30^ and metaFlye^31^ we assembled the long-read (*n*=8 SMRT cells, 74,953 contigs, *N50* = 18.2 kb) into assemblies that totalled 781 Mb in size, with half of the sequence present in contigs longer than 20 kb. The longest contig was more than 753 kb in length assembled from the largest long-read metagenome (Table S2). The two short-read Illumina libraries assembled into ~8 million contigs (3.7 Gb, *N50* = 1 kb).

We also co-assembled the metagenomes to access even more BGC diversity than was permissible from the individual assemblies. We co-assembled the five largest long-read metagenomes which yielded 1.4 Gb of assembled sequence (Table S2) with the longest contig exceeding 1.3 Mb in length (*N50* = 36 kb). This co-assembly was as large as our hybrid co-assembly of two short-read Illumina libraries and four long-read libraries produced with metaSPAdes^32^ (1.7 Gb, *N50* = 2.3 kb). Putative misassemblies identified through MetaQUAST were identified and removed^33^. Overall, the long-read assemblies and co-assemblies produced the largest number of ultra-long contigs (>50 kb) and were thus most suited for the investigation of full-length biosynthetic gene clusters. Together they gave unprecedented access to the BGCs encoded by uncultivated microbes including 1,191 BGCs from the long-read co-assembled metagenome.

Overall, the long-read metagenomes, and particularly their co-assemblies, offered substantially deeper insight into biocrust secondary metabolism than was possible through short-read sequencing and assembly (Fig. S2). For example, the Sequel II assembly had 548 BGCs including 174 full-length BGCs (i.e., the BGC was not truncated on either contig edge), while the short-read assemblies had 359 BGCs between them yet only 9 full-length BGCs. In total, we predict that 712 BGCs are full-length clusters.

The single largest BGC was identified in the ultra-large co-assembly and was putatively assigned to the genus *Nostoc*. It encodes a novel hybrid transAT-PKS-NRPS of 111 kb length, harboring 6 core biosynthetic genes and 8 additional biosynthetic genes. Manual inspection suggests it is full-length, making it one of the longest BGCs to be identified directly from a soil metagenome (Fig. S3). Clearly the co-assembly of multiple long-read metagenomes offers access to a deeper spectrum of BGCs while the diversity of these clusters found here suggests that much secondary metabolic potential remains unrealized in current databases. Moreover, the use of long-read sequencing is central to finding novel full-length gene clusters, an issue that precluded the use of short-read metagenomics previously.

## Thousands of novel gene clusters recovered from biocrust metagenomes

We performed gene cluster identification and annotation for secondary metabolites^34^ using all the *de novo* metagenome assemblies owing to their high contiguity in assembly and high proportion of contigs longer than 5 kb (Fig. 1c). This approach recovered 2,988 biosynthetic gene clusters (BGCs) predicted to produce secondary metabolites from uncultivated biocrust microbes. These span all major secondary metabolite classes with terpenes, ribosomally synthesized and post-translationally modified peptides (RiPPs) and non-ribosomal peptide synthetases (NRPSs) particularly well represented. Cyanobacteria were rich in NRPSs and Type 1 polyketide synthases (PKS) and harbored the most BGCs overall encoding some 1,470 BGCs (Fig. 1d; Table S3). Four hundred and twenty of these non-redundant BGCs could be assigned to the genus *Microcoleus* – the pioneer microbial guild of biocrust^35^. Next, we determined the genetic novelty of our BGCs by evaluating whether previous sequencing efforts had captured the sequence by making queries to the entire NCBI nt sequence database (accessed December 6, 2019^36^). Using thresholds of 75% sequence identity over 80% of the sequence length^37^ we identified 175 BGCs that had been sequenced previously. Thus ~94% of BGCs had not been sequenced before. This reaffirms that biocrust are a rich source of BGCs and underscores the potential for long-read metagenomic sequencing in novel BGCs discovery.

Of the known clusters, 143 belonged to Cyanobacteria including the late branching genera *Microcoleus, Nostoc*, and *Oscillatoria*. BGCs identified from non-cyanobacterial contigs had interesting novel elements. For example, *Planctomycetes* were rich in acyl-amino acids, while *Alphaproteobacteria* had unusually high numbers of the dipeptide N-acetylglutaminylglutamine amides (NAGGN) as well as N-acyl-homoserine lactones that may be involved in quorum sensing^38^. Moreover, many terpenes and Type 3 PKS belonged to the dominant heterotrophic phyla *Alphaproteobacteria* and *Actinobacteria* (Fig. 1*e*). We also found 17 phenazines in our dataset, some of which may have functions in redox balance during anoxia^39^, most of which belonged to cyanobacteria.

## Constitutive transcription of secondary metabolite gene clusters

Desert biocrust communities are sensitive to rain events, as revealed by dramatic changes in microbial community structure^28^ and core gene expression by DNA microarray^22, 40^. To identify secondary metabolite BGCs involved in these dynamics, we mapped 13 biocrust metatranscriptomes to our metagenome assemblies. The metatranscriptomes are from a simulated rain event in the laboratory using intact biocrust from the same site (Moab, UT, USA)^40^. They capture microbial transcription following a wetting event for three diurnal cycles at a resolution of 10 individual timepoints. Like the metagenomic data, 16S rRNA transcript analysis using ESVs from the metatranscriptomic datasets revealed an abundance of transcripts from *Cyanobacteria*, and especially *Microcoleus vaginatus* at all timepoints (Fig. 2*a*). We observed a dramatic increase in 16S rRNA transcript copy numbers across all taxa 15 minutes and 1 hour after wetting possibly indicating increased microbial growth on substrates released during cell membrane permeabilization after wetting^41^ or simply ribosome synthesis as microbes emerge from dormancy.

**Fig 2.**
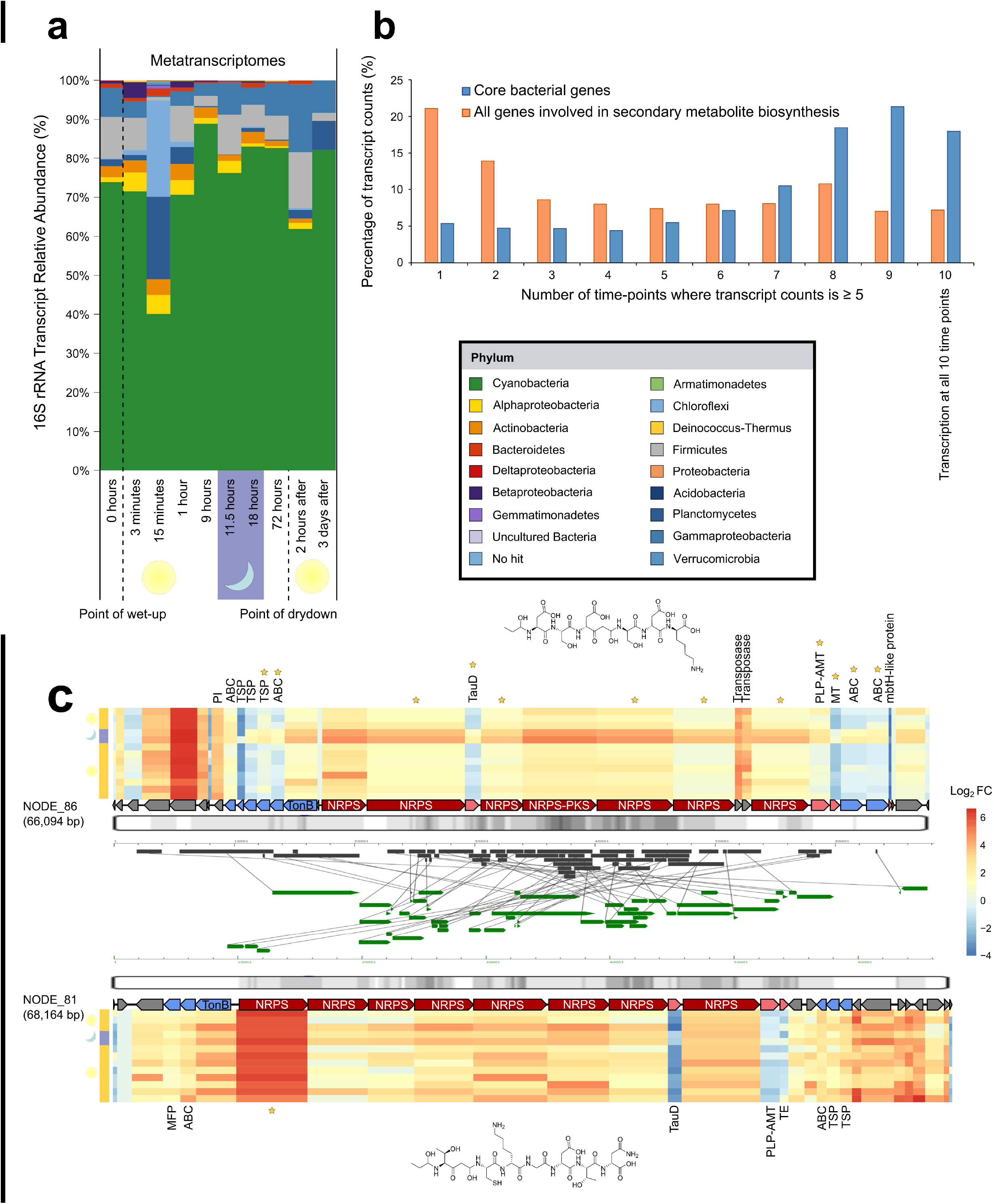
Transcription of secondary metabolites. Transcription of secondary metabolites. **a**, Taxonomic composition of the metagenomes based on exact sequence variants (ESVs) of 16S rRNA transcripts during a soil wetting experiment. Relative abundances were calculated after assigning taxonomy against the SILVA reference database. **b**, Core bacterial gene transcription (*n*=46 genes including DNA-binding proteins, Large and Small subunit ribosomal proteins) shown in blue compared to secondary metabolite gene transcription (orange). Genes transcribed at all 10 timepoints (rightmost point) are thought to experience constitutive expression. The *y*-axis indicates the proportion of genes present in each category. **c**, Putative rearranged siderophore-producing gene clusters found in the co-assembled metagenomes that show homology. Transcriptional profiles of gene clusters with differentially expressed genes. Heatmap columns are scaled to the size of the mapped gene, and row order indicates progression across the experiment from 0 hours (bottom row) to 3 days after wetting (top row). Predicted chemical structures are shown on the right.

The metatranscriptomic data comprised 137 Gb of high-quality sequence in 919 million transcripts from 13 samples (Tables S1, S2, S4). To calculate secondary metabolite gene transcription after wetting we mapped the individual read transcripts to each contig containing a BGC using BBMap^42^ which leveraged our long contigs to profile transcription for almost 3,000 secondary metabolite gene clusters. Remarkably, we found that 395 genes from 240 BGCs were transcribed at all timepoints (using a threshold of at least five mapped transcripts per gene within a cluster at a single time point), which represent some 6% of all secondary metabolic genes in our dataset (Fig. 2*b*). Our results show stark contrast to previous observations that BGC expression in the laboratory is low wherein most secondary metabolites are not transcribed^2^. Their constitutive expression supports the notion that “secondary” metabolites may play critical (and possibly essential) roles in communication or niche occupancy in these ecosystems. Given the relatively high biosynthetic cost of synthesizing secondary metabolites *vs*. primary metabolites^43^ this suggests that these compounds provide fitness benefits to their hosts across the wetting event.

Next, we investigated how the observed constitutive expression of secondary metabolic genes compared to the transcription of all other genes, i.e., those not involved in secondary metabolism. Of these 966,111 ‘non-secondary’ genes, just 43,139 (some ~4.5%) were constitutively transcribed at all 10 time points. These mapping rates were not artifacts of gene length differences between primary genes and secondary metabolic gene lengths (Supplementary Results and Fig. S4). We then focused on core 46 metabolic bacterial genes that we expect to have high constitutive expression e.g., those encoding DNA-binding or ribosomal subunit proteins (Table S5), and found that indeed many of these core genes were transcribed at eight or more time points and 18% that were constitutively transcribed (5 mapped transcripts at all 10 timepoints; Fig. 2*b*). This same analysis of secondary metabolic genes showed a more even distribution across the time points with 6% transcribed at all 10 time points (Fig. 2*b*). Although lower than for core bacterial genes, this represents a higher proportion of constitutive transcription for secondary metabolic genes than was anticipated.

While our results show low level constitutive transcription of many BGCs, the highest level of BGC transcriptional activity occurred at night, 11.5 hours after the initial wetting event (Fig. S5*a*). This enrichment in transcription was mostly underpinned by a surge in transcriptional activity by the *Cyanobacteria* (Fig. S5*b*) which likely corresponds to gene induction at night when they are not photosynthetically active^40^. Strikingly, 80% of cyanobacterial BGC transcription peaked at night (2,527 of 3,173 genes). This included the significant transcription of two putative siderophore-producing BGCs (DESeq2: *P* < 0.05), while their observed rearrangements were presumably driven by transposases (Fig. 2*c*, Fig. S8 and Supplementary Results).

We next examined phylogenetic conservation of BGC expression among phyla. Here, we analysed a subset of biosynthetic genes individually (*n*=12,470 genes) using t-SNE visualization^44^. This analysis revealed segregation of biosynthetic gene transcription by taxonomy (Fig. 3*a*). Cyanobacterial transcription of secondary metabolites was significantly unlike all other phyla (Pearson’s *r* > 0.8; adjusted *P* < 0.05). While Cyanobacteria exhibited the highest level of BGCs transcription at night, 11.5 hours after wetting, other bacteria (in this case almost exclusively heterotrophic guilds) showed maximal BGC transcription during the day (Fig. S5). Notably there was a peak of transcriptional activity 72 hours after wetting (during the day, and the point of dry down) which was due to the increased transcription of terpenes and Type3 PKSs by abundant heterotrophic bacteria such as *Deltaproteobacteria* and *Actinobacteria* (Fig. S6, S7*b*).

**Fig 3.**
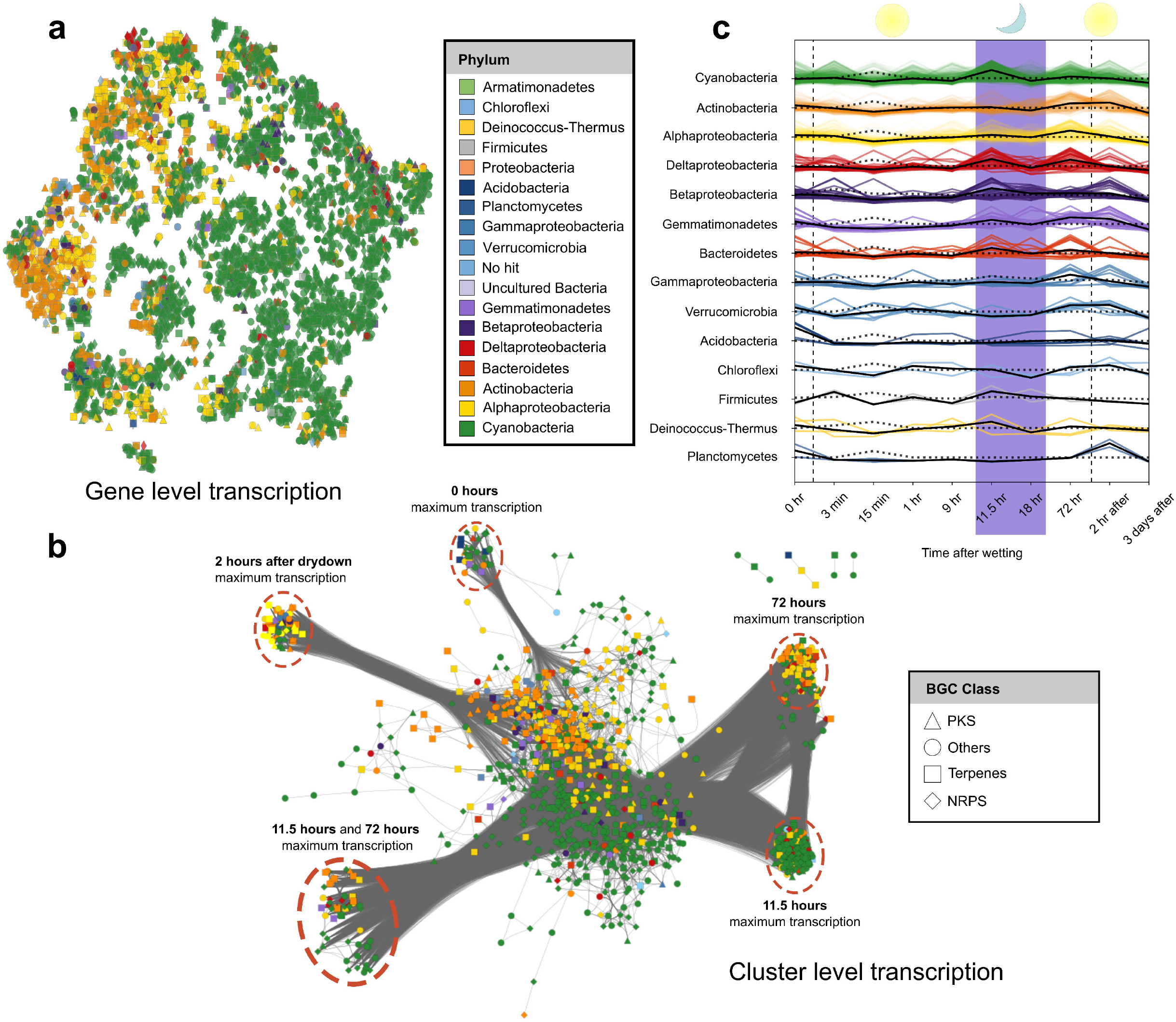
Phylum specific transcription of secondary metabolites. **a**, t-SNE visualization of every individual biosynthetic gene identified. The color of the points indicate the phylum assignment whilst shapes indicate the BGC class. **b**, Co-occurrence network based on Pearson correlations (*r* > 0.8) among entire BGCs (*n*=2,988) based on average z-scores at each time point. Each node is a BGC within a contig that are colored by phylum and shaped by BGC type. Closely clustered nodes share similar transcriptional profiles. **c**, Line plot showing 16S rRNA transcript copy number over time shown by black, dotted lines. Average BGC transcription over time shown by the colored, solid lines. *Cyanobacteria* (green) show a unique night-time upregulation of secondary metabolism. Purple background indicates night-time transcription.

Given the conserved phylogenetic signal level at the gene level, we also examined phylogenetic conservation at the cluster level. Here we compare the degrees to which biosynthetic gene clusters shared similar transcriptional profiles across phyla using a co-occurrence network based on the average Z-scores of each BGCs transcription (*n*=2,988). This analysis revealed clustering of secondary metabolite transcription of entire BGCs by taxonomy. Namely, the bacterial phyla had distinct temporal signatures of BGC transcription compared to each other over the course of 3 days (Fig. 3*b*). Cyanobacterial BGC expression was distinct from all other bacterial groups in the biocrust (Fig. 3*c*; *P* < 0.05). To our knowledge, this is the first such observation of phylum-level differences in microbial BGC transcription in natural communities. This may reflect conservation of life history traits especially niche competition strategies. For example cyanobacteria can grow heterotrophically on diverse dissolved organic components^45^ and increased BGC expression may reflect increased competition with heterotrophs occurring at night. Thus, at night *Microcoleus* and other cyanobacteria may produce antibiotics to antagonize heterotrophs competing for dissolved organic compounds^46^.

In addition to antagonism, night-time expression of BGC products can facilitate electron and nutrient transport. Redox-active secondary metabolites are known to be produced by microbes under anoxic conditions^39^. For example, *Pseudomonas aeruginosa* enhances substrate-level phosphorylation during anoxia through the production of phenazines that facilitate electron transport^47^. Constitutive expression of the siderophore-producing gene clusters in cyanobacteria may reflect cation import strategies (notably iron scavenging) needed to support photosynthesis and other metabolic activities (Supplementary Results).

## Conclusion

In this study we show that long-read metagenomic sequencing is a powerful new tool for the examination of secondary metabolite gene clusters directly from complex environmental samples. Integration with metatranscriptomics revealed that ~6% of secondary metabolic genes were constitutively transcribed over 3 days – a higher percentage than other genes. Thus, while conventionally unexpressed under laboratory conditions, our results show that *in situ* BGCs appear to control important life history traits involved in maintaining microbial niches. BGC expression showed strong phylogenetic conservation where *Cyanobacteria*, unlike other phyla, exhibited the highest levels of transcription at night. We speculate that this may reflect the switch from cyanobacteria serving as primary producers during the day to competing with heterotrophs for dissolved organics at night.

## Materials and Methods

### Biocrust Sample Collection and DNA isolation

Biological soil crust (biocrust) was collected from Green Butte Site near Moab, UT, USA (38°42’54.1’’N, 109°41’27.0”W) in 2014 as described previously^22^. This field site is part of a long-term ecological research area of scientific interest aimed at exploring climatic changes in arid regions. We sampled early maturity biocrust (*Microcoleus*-dominated) by coring directly into the soil surface with a petri dish (6 cm^2^ by 1 cm in depth). Samples were maintained in petri dishes in a dark desiccator in the laboratory until required for DNA isolation. Metagenomic DNA was isolated using the MoBio Powersoil kit as per the manufacturer’s instructions with a minor modification. We extracted DNA from 2 g of crust material by dividing the sample into four separate tubes (0.5 g in each tube). The nucleic acids from each tube were eluted in 50 μl of elution buffer and then pooled these into a final sample containing 200 μl of elution buffer and DNA.

### SMRT Sequencing

We sequenced three SMRT cells on the PacBio RS II Single Molecule, Real-Time (SMRT®) DNA Sequencing System (Pacific Biosciences, CA, USA) using two different library inserts: 10 kb AMPure PB library [*n*=2] and a Low input 3 kb PB library [*n*=1] using binding kit P6 v2 with 360-minute and 120-minute movies for the respective libraries. The same libraries were then sequenced on a PacBio Sequel System (Pacific Biosciences) using Sequel Binding Kit 2.1 with a combination of 600- and 1200-minute movies. A third library was made using 10 kb AMPure PB approach with a Blue Pippin size cutoff of 4.5 kb. It was sequenced on PacBio Sequel II System (Pacific Biosciences) using 1.0 template prep kit and a 900-minute movie.

To test how well-suited long-read metagenomes are for BGC recovery, we made use of five publicly available PacBio SMRT metagenomes including a biogas reactor library sequenced on the PacBio RS II System with a 2 kb insert length^48^, and four metagenomes obtained from Lake Biwa, Japan that were sequenced on a PacBio Sequel System with a 4 kb insertion length^49^. Raw sequence statistics for each metagenome is provided in Table S1. We analysed the sequencing effort of the metagenomes using Nonpareil v3.30^50^ which relies on read redundancy. We performed a similar comparison using publicly-available long-read metagenomes which also yielded improvements in contig sizes and BGC yield from co-assembled datasets (Supplementary Material).

### Illumina Sequencing

Two unamplified 300 bp Illumina libraries were generated and sequenced 2×150 bp on the HiSeq-2500 1TB platform (Illumina).

### Taxonomy

We extracted prokaryotic 16S rRNA genes using SortMeRNA 2.1b^51^. These 16S rRNA sequences were then analysed using DADA2^52^ to identify exact sequence variants (ESVs) under default parameters with the exceptions of truncLen (150) and maxEE (1). The ESVs were then assigned taxonomy against the entire SILVA 16S rRNA gene reference database^53^. The taxonomy of the identified gene clusters was inferred by BLAST queries^36^ against the NCBI nr-database whereby hits were retained with E-values of less than 1 × 10^-10^ and bit scores greater than 60.

### Assembly

We performed read correction, trimming and assembly for the three RS II SMRT cells with Canu v1.8^30^. Here we included parameters suggested by the developers of Canu for PacBio metagenomes including an estimated mean genome size of 5 Mb (genomeSize=5m). We also changed the following parameters from their default values: corMinCoverage=0, corOutCoverage=all, corMhapSensitivity=high, correctedErrorRate=0.105, corMaxEvidenceCoverageLocal=10 and corMaxEvidenceCoverageGlobal=10.

The four larger Sequel metagenomes were assembled using metaFlye v2.4.2 under default settings with an estimated genome size of 5 Mb and the –*meta* option implemented for metagenomic sequence data^31^. All Illumina sequence data were quality trimmed prior to assembly using Prinseq-lite v0.20.4^54^ with --*min_qual_mean* set to 20 and -*ns_max_n* set to 0 which eliminates low quality reads and ambiguous bases (internal N’s). We assembled the two biocrust Illumina metagenomes with metaSPAdes v3.13.0^32^ as recommended for paired-end short read length Illumina libraries^55^. We also co-assembled the four Sequel libraries together (termed Flye co-assembly), and then with the Sequel II library (termed Ultimate co-assembly) using metaFlye. Finally, we co-assembled the four Sequel libraries with the two Illumina metagenomes using metaSPAdes. Open reading frames (ORFs) of core metabolic genes were predicted from the assembled metagenomes using Prodigal^56^ and annotated using Prokka^57^ in KBase (https://kbase.us/)^58^. All assemblies were quality-checked using MetaQUAST^33^ which precluded the inclusion of misassemblies from our analysis.

### Biosynthetic Gene Cluster Analysis

All contigs > 5 kb in length were explored for biosynthetic gene clusters (BGCs) using the antiSMASH v5.0 web server under strict settings^34^. Next, we consolidated and passed all putative BGCs through BiG-SCAPE v0.0.0r and CORASON in glocal mode to explore the phylogenomic relationships between the BGCs recovered from the 11 biocrust metagenomic datasets^13^. BiG-SCAPE consolidates both antiSMASH and the MiBIG 2.0 database to support initial antiSMASH predictions and so we included the entire MiBIG 2.0 database in our analysis to place our BGCs among verified clusters^59^.

To determine the genetic novelty of our BGCs we performed homology searches against the NCBI nt database (downloaded December 6th, 2019) using NCBI blast+ 2.9. We only retained top hits based on an E-value of 1 × 10^-10^. BGCs were non-redundant (not sequenced previously and thus novel) if sequences matched ≤ 80% of the BGC query length and had an average of ≤ 75% sequence identity against the database. We corroborated the taxonomic assignments using the Contig Annotation Tool under default settings (CAT, v5.0.4)^60^. Chemical structure predictions were first created by antiSMASH v5.0.

### Metatranscriptomic mapping

We made use of metatranscriptomes sequenced from biocrust material collected at the same sampling site in Moab, Utah that were publicly-available on JGI GOLD^40^. The experimental design tracked the transcriptional responses of biocrust communities over two complete diurnal cycles following an artificial wetting event in the laboratory with 12 hours of light followed by 12 hours of dark (Table S1). The time points at which transcripts were collected include: 0 hours (immediately before wet-up), 3 min, 15 min, 1 hour, 9 hours, 11.5 hours, and 18 hours after wet up, then 72 hours after wet up (immediately prior to dry down), then 2 hours and 3 days after dry down. The 11.5 hours and 18 hours samples also represent transcriptional activity at night-time while all other samples captured transcription during the day.

Transcripts were quality-controlled using Prinseq-lite v0.20.4 as described above for the Illumina data. The metatranscriptomes were then assembled using metaSPAdes. The unassembled transcripts were then mapped to contigs containing BGCs using bbmap v38.73^42^. We used SAMtools v1.9^61^ for file conversion and sorting. Mapped sequences and associated contigs were then visualized within Geneious^62^. We used DESeq2 v1.28.0^63^ in the R statistical environment v3.6.3 to test which genes underwent differential expression by explicitly testing expression against the control sample (0 hours). Here we tested two environmental treatments, (i) the diurnal cycling regime (i.e., day to night to day) and, independently, (ii) the influence of wetting and drying. Transcripts were removed that did not map at the phylum level to the 16S data or that had a maximum count less than 20 in any sample. The remaining transcript levels were normalized by the total counts for each sample and then multiplied by the average count across all samples. Duplicate samples at the 15-minute time point and triplicate samples at the 1-hour timepoint were averaged, and z-scores of normalized transcript abundance mapped to each biosynthetic gene to reveal which time points showed highest gene activity. In addition, z-scores were used with t-SNE (T-distributed Stochastic Neighbor Embedding) to visualize the gene transcription patterns in ordinance space^44^. The t-SNE implementation in sklearn (v 0.23.2) manifold module was used with the following parameters: ‘angle’: 0.5, ‘early_exaggeration’: 12.0, ‘init’: ‘random’, ‘learning_rate’: 200.0, ‘method’: ‘barnes_hut’, ‘metric’: ‘euclidean’, ‘min_grad_norm’: 1e-07, ‘n_components’: 2, ‘n_ite?: 3000, ‘n_iter_without_progress’: 300, ‘perplexity’: 40, ‘random_state’: None, ‘verbose’: 1.

## Supporting information

Supplementary Results and Figures

## Acknowledgements

This work was partially supported by funds provided by the Office of Science Early Career Research Program Office of Biological and Environmental Research, of the U.S. Department of Energy and by the U.S. Department of Energy Joint Genome Institute, a DOE Office of Science User Facility, supported by the Office of Science of the U.S. Department of Energy under Contract No. DE-AC02-05CH11231 to Lawrence Berkeley National Laboratory. We also wish to acknowledge Simon Roux, Emiley Eloe-Fadrosh and Eoin Brodie for their constructive feedback.

## Author contributions

M.W.V.G. conducted the metagenomic analyses, formulated ideas, and wrote the manuscript. A.R.O. predicted biosynthetic gene clusters, performed DESeq2 analysis and wrote the manuscript. B.P.B. conducted the statistical analyses among transcriptional profiles and produced figures. P.F.A. assisted with contig annotation and provided ideas. T.L.S. collected samples and extracted DNA. A.C. provided access to data, collaborated with Pacific Biosciences scientists to produce long-read metagenomes and offered valuable insights into the sequence data. R.R. assembled the large metagenomes and assisted with transcript mapping. G.H. developed technology of high-molecular weight DNA shearing and size selection optimization for library construction in collaboration with Pacific Biosciences scientists. M.K. developed Pacific Biosciences SMRTbell template preparation methods for HiFi 10 kb libraries from sample quality assessment to library preparation including development of multiplex strategy. L.S. and M. Y. optimized sequencing conditions for metagenomic sample libraries on the Pacific Biosciences RS II, Sequel and Sequel II sequencer platforms in collaboration with Pacific Biosciences. C.G.D. contributed sequencing platform development and operation management to enable the sequencing of environmental metagenomic samples on the Pacific Biosciences sequencing instruments in collaboration with Pacific Biosciences. Y.Y. developed long-read technology for environmental samples from design/coordination to execution in collaboration with Pacific Biosciences. T.P.M. offered logistical support and constructive feedback on the manuscript. F.G.P. provided access to Microcoleus genomes and assisted with meaningful interpretation of the cyanobacterial transcriptomics data. R.C.O. contributed overseeing the development of long-read technologies including 10kb HiFi sequencing for environmental metagenomics in collaboration with Pacific Biosciences and JGI staff. A.V. wrote the paper and offered substantial advice on project planning. L.A.P. assisted in the interpretation of biological and genomic data. T.R.N. Designed the study and wrote the manuscript.

## Data availability

Raw data of the long- and short-read biocrust metagenomes can be accessed on the IMG/M website (Submission ID 241874) or on the NCBI website (BioProject: PRNJNA691698). The raw metatranscriptomic data are publicly-available through the JGI GOLD portal (sequence project IDs 1010318 – 1022409).

