## Supplementary Results and Figures for "Long-read metagenomics of soil communities reveals phylum-specific secondary metabolite dynamics"

Prepared for: *Nature Microbiology*

**Supplementary Methods**

**Publicly-available long-read metagenomes**

We downloaded four Sequel metagenomes (Pacific Biosciences) derived from Lake Biwa, Japan ^1^ as well as RS II and Illumina metagenomes from a biogas reactor ^2^ (Table S1). All PacBio metagenomes were assembled with Canu v1.8 ^3^ using the following parameters: genome size of 5 Mb (*genomeSize*=5m), *corMinCoverage*=0, *corOutCoverage*=all, *corMhapSensitivity*=high, *correctedErrorRate*=0.105, *corMaxEvidenceCoverageLocal*=10, *corMaxEvidenceCoverageGlobal*=10. metaSPAdes ^4^ was used for the short-read biogas metagenome and the co-assembly of both biogas metagenomes. Unfortunately, we were unable to assemble the biogas PacBio metagenome using either Canu or metaFlye ^5^, possibly owing to insufficient sequence coverage to generate contigs. We also co-assembled the four Lake Biwa Sequel metagenomes using Canu. The quality of each assembly was provided by MetaQUAST v4.6.3 ^6^ and is available in Table S2. We then predicted BGCs from the assembled long-read metagenomes using antiSMASH v5.0 ^7^.

The results of this analysis showed that, as for the biocrust metagenome assemblies, that co-assembling multiple metagenomes dramatically improves yield of BGCs. For example, the Lake Biwa samples produced 50 BGCs individually, but when co-assembled yielded 59 BGCs. This modest improvement was also observed in full-length BGCs (16 compared to 18).

**Supplementary Results and Discussion**

**Expression of Cyanobacterial Siderophore BGCs at Night**

Differential gene expression analysis (DESeq2 ^8^; *P* < 0.05; FDR = 5%, Table S6) using mapped transcripts revealed that 10 BGCs contained biosynthetic genes that underwent significantly more transcription at night, all of which were cyanobacterial in origin. The most dramatic of these enrichments involved two cyanobacterial NRPS-PKS hybrid BGCs, identified on Node_81 and Node_86. These BGCs likely encode for a novel siderophore, putatively assigned based on genes encoding predicted membrane proteins involved in siderophore and iron transport located in the clusters (Fig. 2*c*). Additionally, a subset of cation acquisition genes was upregulated at night, suggesting a multifaceted approach for cation import at night that is under significant control of native regulatory constraints.

Node_81 (68 kb), appears to form a novel heptapeptide, while Node_86 (66 kb) is 2 kb shorter and has one fewer NRPS module, thus forming a hexapeptide (Fig. 2*c*). We speculated these to be rearranged BGCs based on the presence of transposases within Node_86, which were supported by differences in G+C content flanking the transposases, potentially indicating recent transposition. Overall, 80% of transposases located in BGCs were active across all time points.

**Cation acquisition at night**

To further investigate the transcriptional activity of cation acquisition genes at night, we mapped reads to all the co-assembled metagenomes. A subset of putative cation acquisition and sequestration gene were differentially expressed, specifically *hemH* (Ferrochelatase), *hxuB* (Heme/hemopexin transporter protein), *pacS* (putative copper-transporting ATPase) and *idiA* (iron ABC transporter) genes (Table S7). The differential expression of siderophore and cation acquisition genes at night suggests a common ‘night-time’ strategy of cation import in biocrust, which was consistent across most cyanobacterial BGCs (Figs. S6, S7).

**Transposases are prevalent among biocrust BGCs**

Two transposases and a phage integrase were found on Node_86, with the transposases located upstream of the final NRPS gene in the cluster. G+C content surrounding the transposases within the BGC had high levels of G-C skew (>1.5% deviation from the mean) potentially indicative of a recent gene rearrangement. We hypothesized that these mobile elements were responsible for the rearrangement of Node_81 into Node_86. This is supported by the perpetual expression of the two transposases and phage integrase in Node_86, implicating a mechanism for BGC rearrangement throughout the experiment and suggesting recombination events may still be occurring. We speculate that BGC re-arrangements may be on-going, long-term processes rather than brief, one-time events. Overall, 20% of transposases (Fig. S8) within BGCs were constitutively transcribed, hinting that similar long-term recombination events may be occurring in the biocrust community. Transposases located outside of BGCs, i.e., those more relevant to primary metabolism, showed less constitutive transcription (~7% of transposases) compared to those involved in secondary metabolism. Moreover, 26% of non-BGC transposases were never transcribed while only 19% of BGC transposases were never transcribed.

Nine of the 10 genes comprising Node_86 showed differential expression, with significantly higher transcription at night. As a putative siderophore, the function of this metabolite could be to acquire iron at night in preparation for photosynthesis the following day. In contrast, Node_81 only had one differentially expressed gene, but still tended towards nighttime activation.

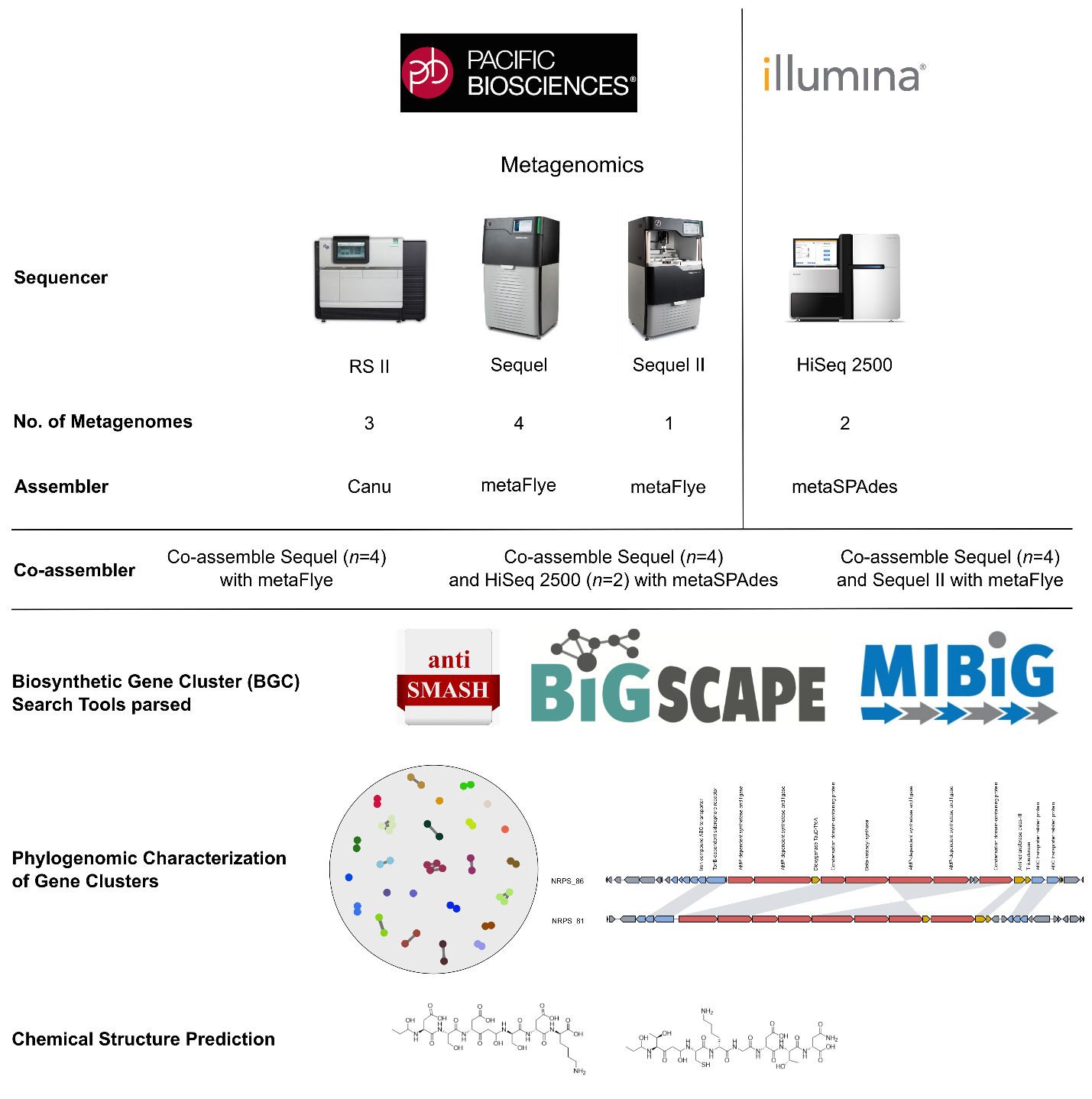
**Fig. S1 | Workflow overview.** 8 long-read metagenomes were generated using Pacific Biosciences instruments that span 3 generations (RS II -> Sequel -> Sequel II). 2 short-read metagenomes were generated from the same biocrust samples using an Illumina HiSeq 2500 instrument.

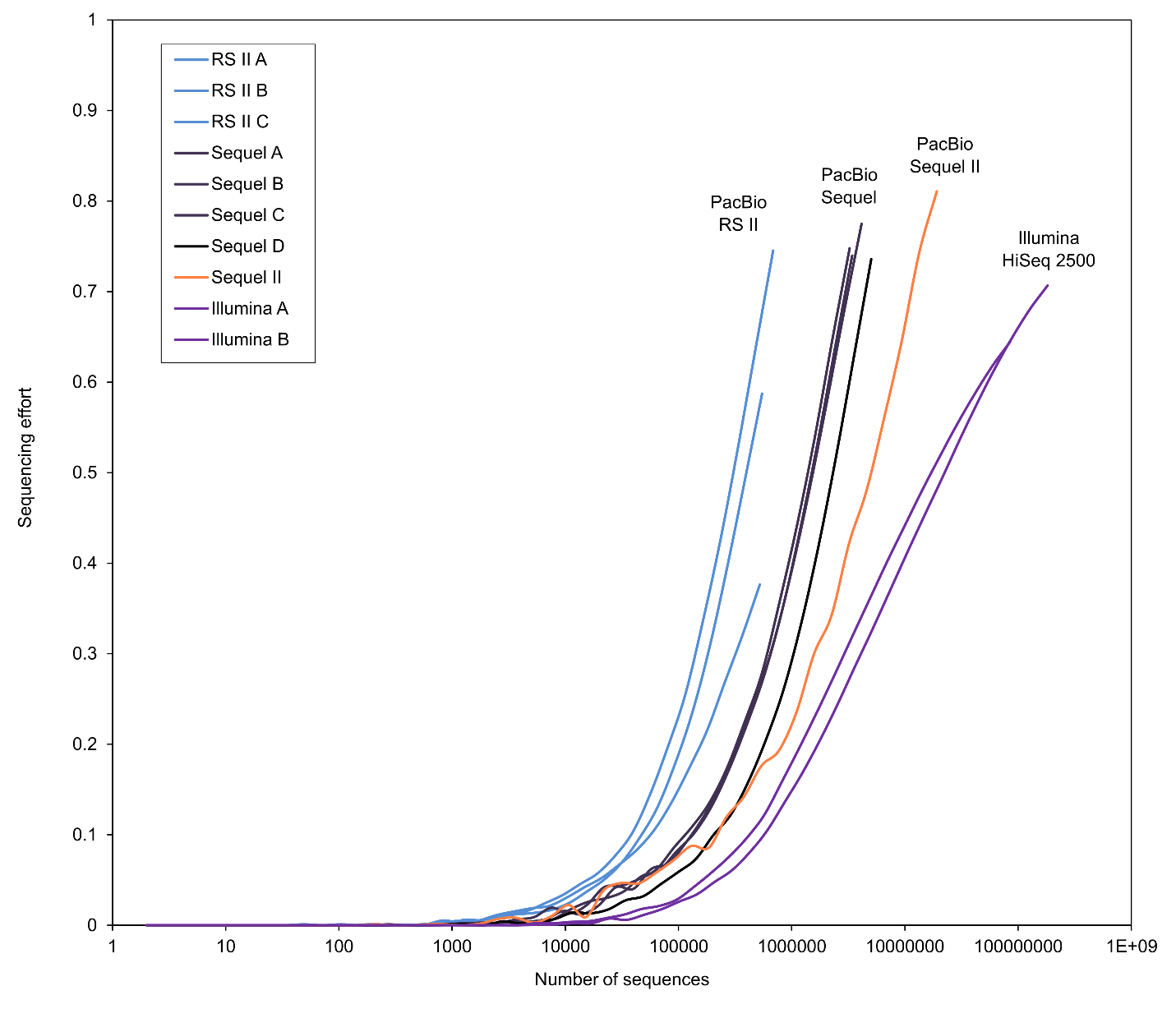
**Fig. S2. | Estimates of sequencing effort.** Nonpareil estimates sequence coverage by calculating read redundancy. The generations of long-read technology (RS II -> Sequel -> Sequel II) lead to improvements in sequencing depth. Short-read sequencing required orders of magnitude more sequencing to achieve a similar sequencing effort as RS II.

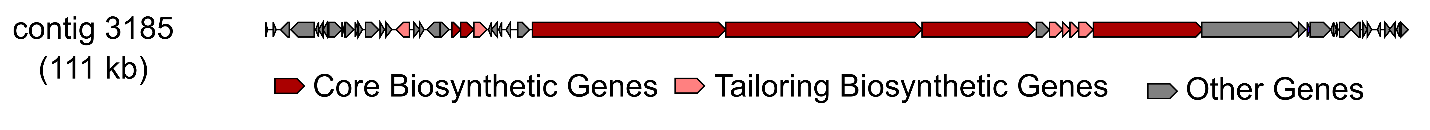

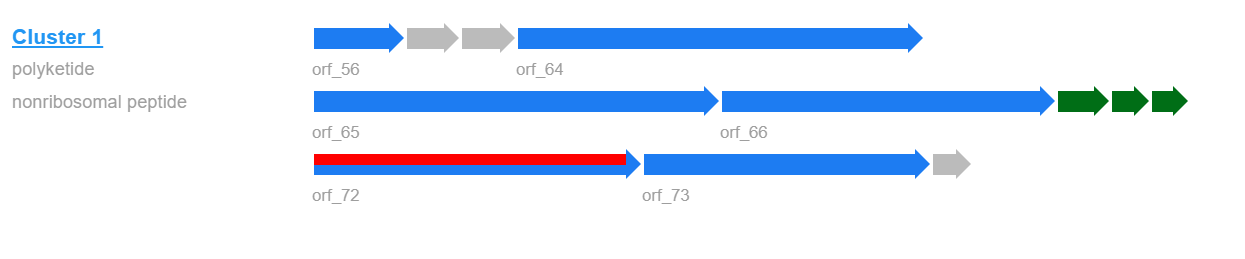

**Figure S3. | Longest BGC recovered from a metagenome.** The longest BGC found across the metagenomes encodes a 111 kb transAT-PKS-NRPS. The domain architecture is provided by PRISM.

**Fig. S4. | Mapping transcripts as a function of gene length.** To test whether gene length influenced mapping rates we compared how read recruitment differed for the longer secondary metabolite genes (average gen e length=1153 bp) compared to the primary metabolic genes (average gene length=688 bp). Visualizing the number of mapped transcripts by gene length showed no correlation for either **a)** secondary or **b)** primary metabolic genes. Similarly, comparing the number of timepoints with 5 or more transcripts (where 0 means never expressed and 10 means constitutive expression) to gene length indicated no trend that followed an increase in transcript recruitment onto longer genes for either **c)** secondary or **d)** primary metabolic genes.

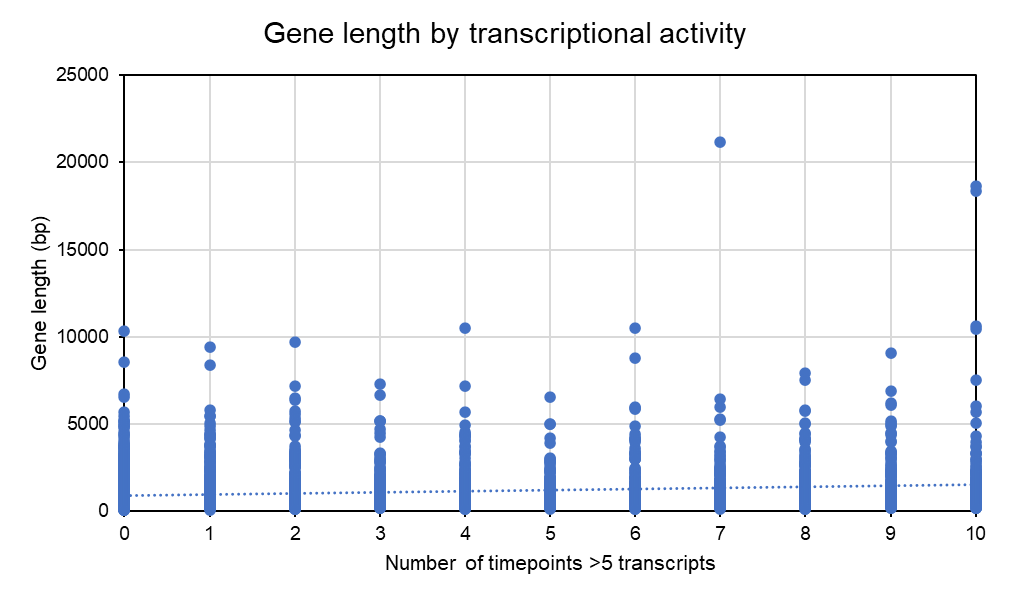

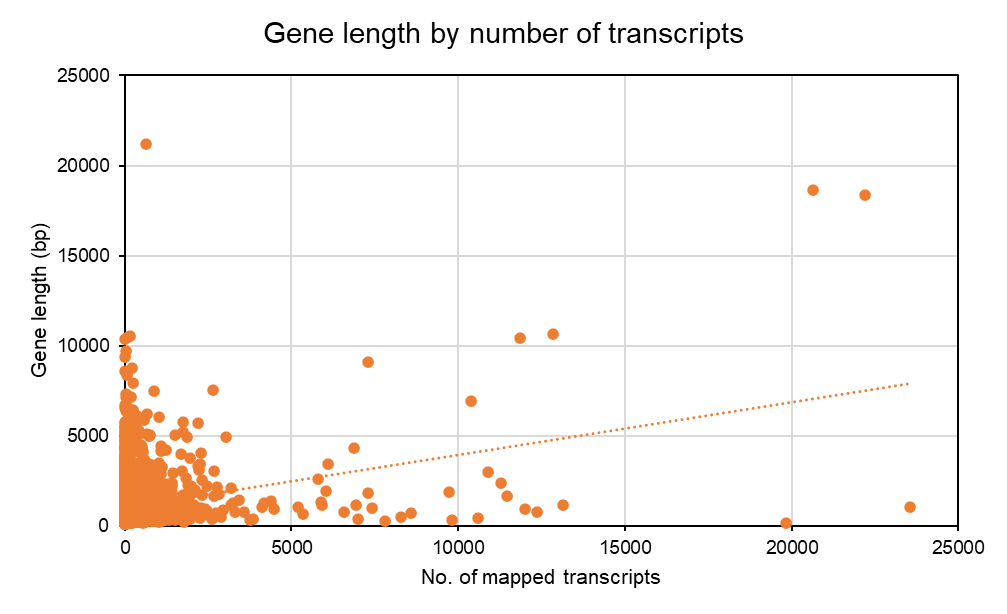

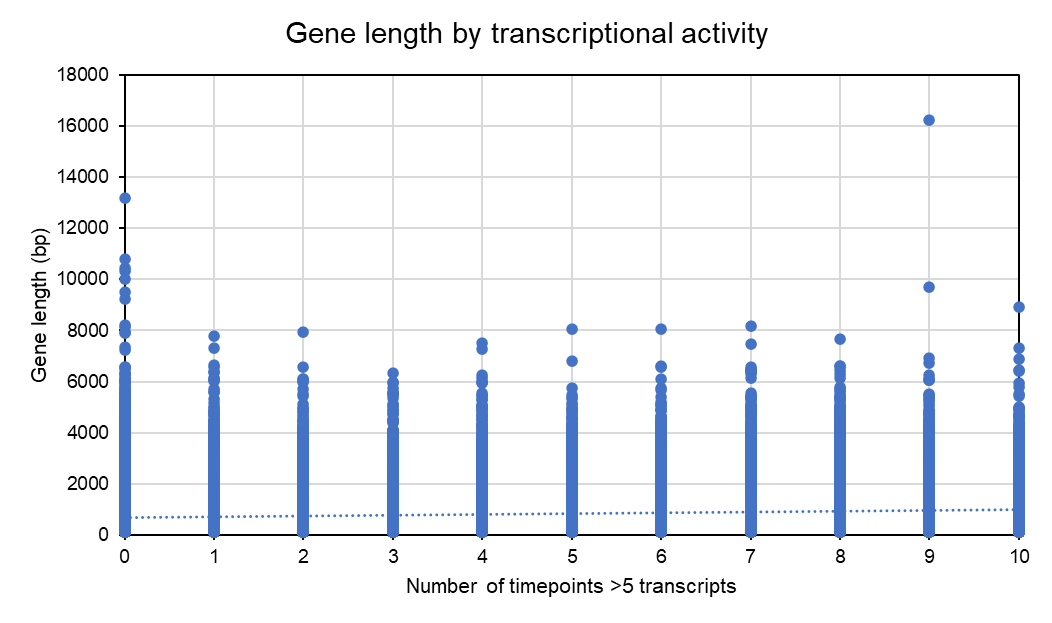

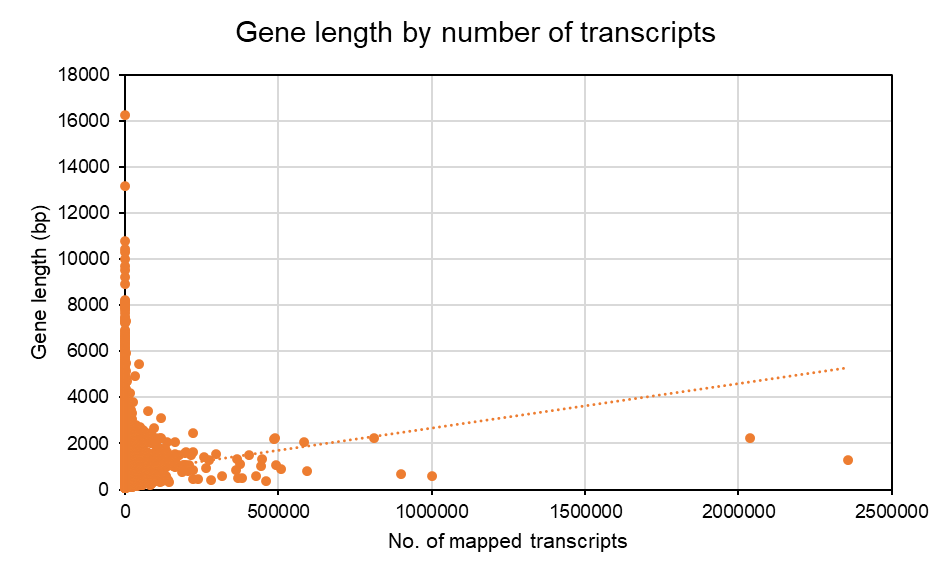

**A**

**B**

**C**

**D**

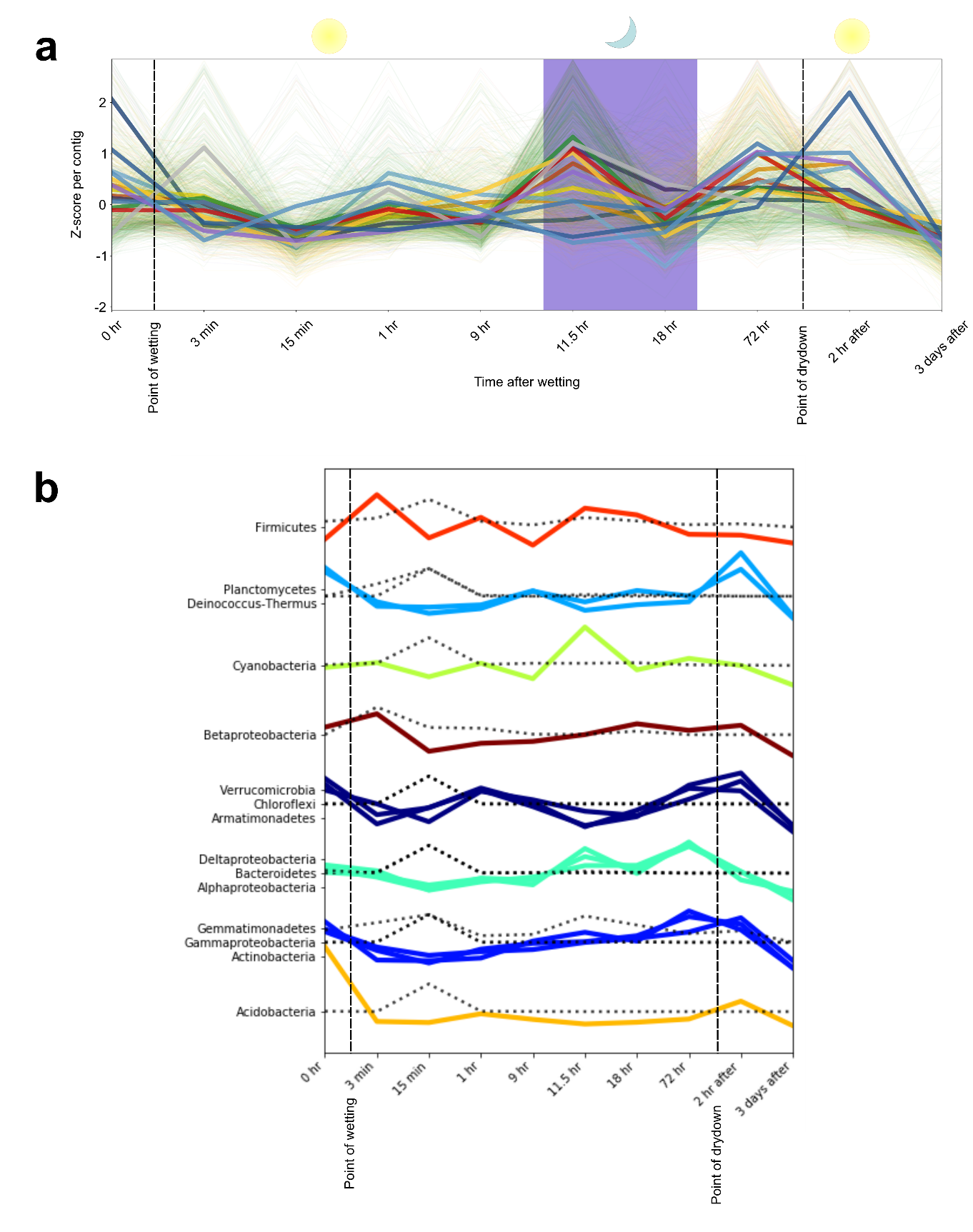
**Fig. S5.** **| Diurnal trends in BGCs.** **a,** Trends across the 3-day phase were detected in 12,470 expressed biosynthetic genes (cutoff > 20 mapped transcripts across time points). Read counts of transcript relative abundances per gene were Z‐score normalized for the purpose of visualization. Each gene trend is color‐coded by its taxonomic affiliation. The purple background indicates night-time transcription. **b,** Clusters of bacterial phyla based on their average Z-score from all contigs with BGCs. The single-colored lines indicate secondary metabolism over time, while the dotted lines indicate the number of 16S rRNA transcripts at each timepoint.

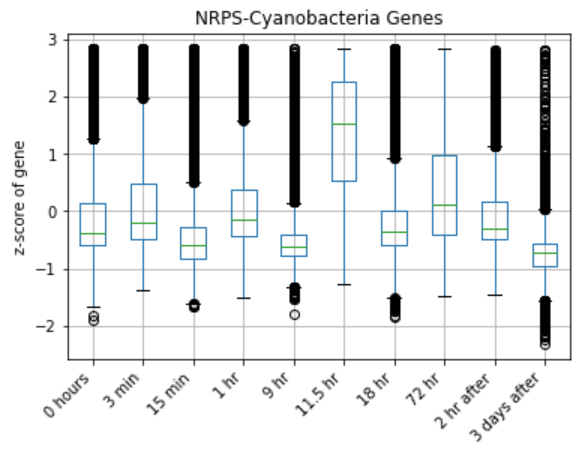

**Fig. S6. | Cyanobacterial transcription occurs primarily at night.** All NRPS gene clusters belonging to *Cyanobacteria* showed significantly more transcriptional activity 11.5 hours after wetting (*P* < 0.05). This first dark time point indicates a shunt of secondary metabolism by *Cyanobacteria* that also includes the increased transcription of RiPPs, T1PKS and T3PKS gene clusters.

**Fig. S7.** | **DESeq2 showed significantly enriched gene transcription at night.** Genes labelled on the heatmaps were those located within BGCs. **a,** Flye co-assembly. **b,** Ultimate co-assembly.

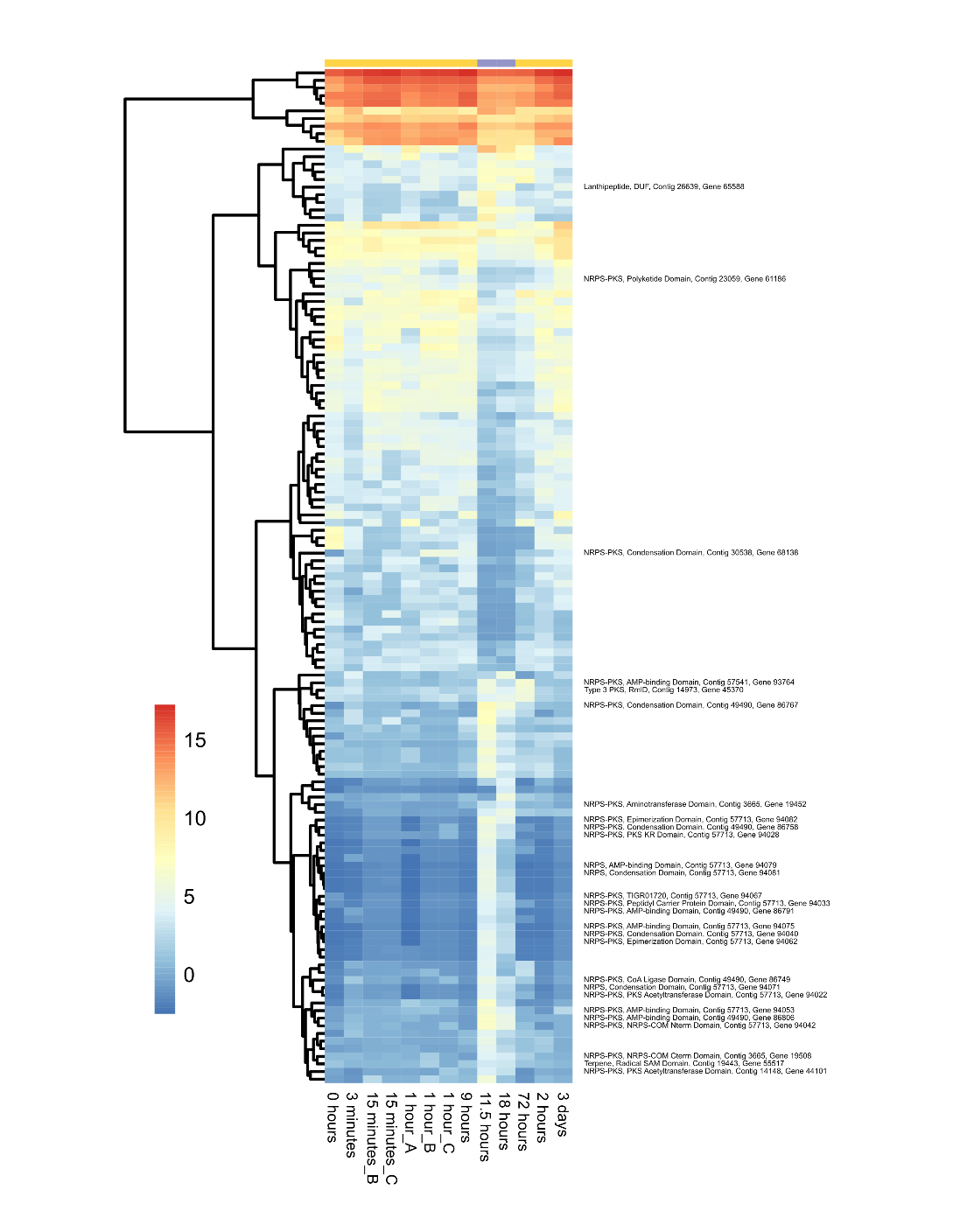

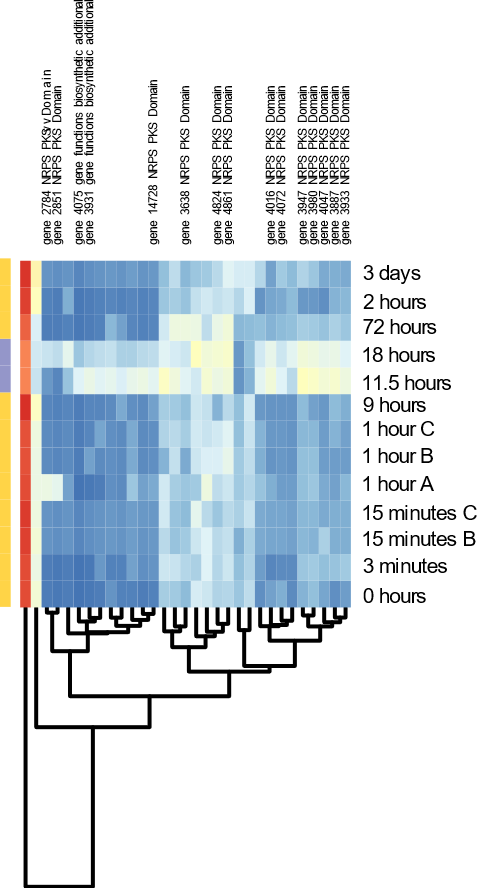

**Fig. S8. | Transposase transcription.** Heatmap showing the transcription of all transposases located in BGCs over time. We identified 3 broad categories of expression: (i) unexpressed (no mapped transcripts), (ii) weak- to moderate-expression, and (iii) strongly expressed transposases.

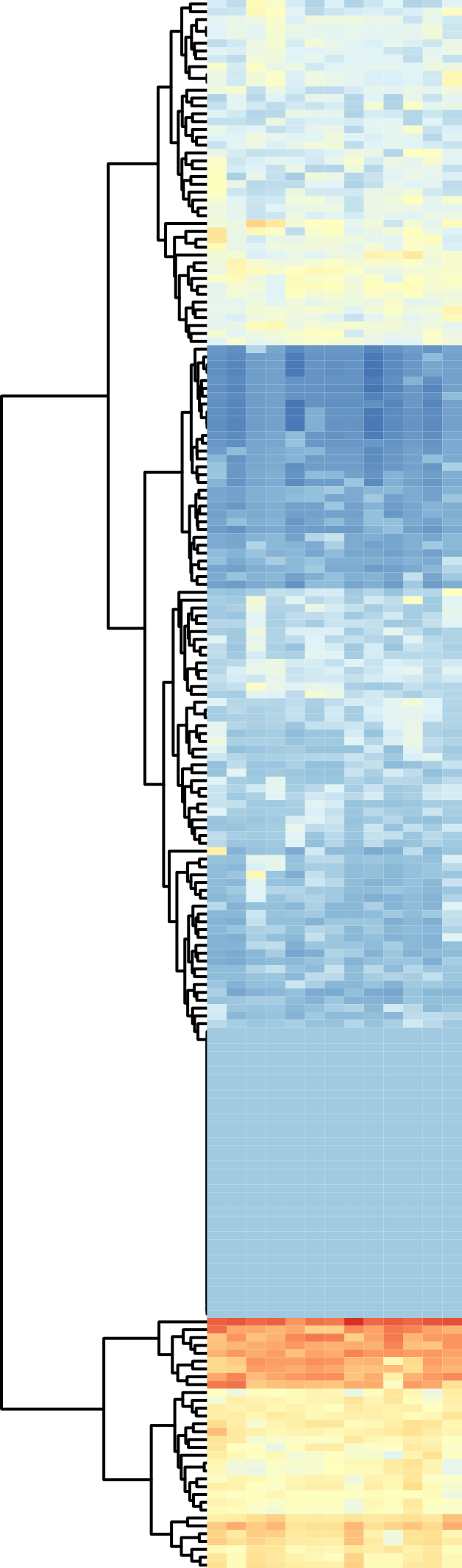

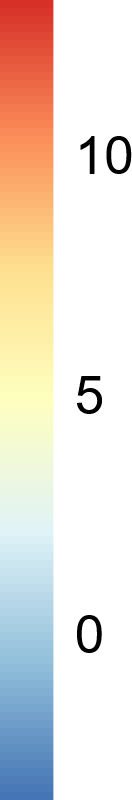

37 unexpressed transposases

32 strongly expressed transposases

131 weakly- to moderately-expressed transposases
